# Genetic Variability and Population Structure within the *Anopheles tessellatus* complex (Theobald, 1901) in Indonesia using ITS2 nuclear and COI, COII mitochondrial sequences

**DOI:** 10.64898/2026.04.08.717322

**Authors:** Anis Nurwidayati, Hari Purwanto, Raden Roro Upiek Ngesti Wibawaning Astuti, Yudhi Ratna Nugraheni, Lulus Susanti, Yuyun Srikandi, Budi Setiadi Daryono, Triwibowo Ambar Garjito, Sylvie Manguin

## Abstract

Some *Anopheles* species that act as malaria vectors are members of species complexes, a concept whereby sibling species cannot be differentiated solely on the basis of morphological characters. Therefore, species complexes represent a major problem in malaria vector control, because within an *Anopheles* complex, vectors cannot be differentiated from non-vector species, unless molecular techniques are used to identify them. The *Anopheles tessellatus* species complex is an important potential vector in South, East, and Southeast Asia, including certain regions of Indonesia. However, no in-depth studies have been conducted on this species complex in that country. Therefore, this study investigated the taxonomic status of *An. tessellatus* from diverse populations across five Indonesian islands (Sumatra, Java, West Nusa Tenggara, East Nusa Tenggara, and Sulawesi) and identified interpopulation genetic variation based on molecular data of the ITS2, COI, and COII genes. Phylogenetic relationships were constructed using the Maximum Likelihood method. Haplotype and network analysis were also conducted. The results indicate that *An. tessellatus* constitutes a monophyletic group comprising three well-defined lineages that exhibit clear intraspecific genetic differentiation. Cluster 1 corresponds to the population of Sumatra, Cluster 2 represents population from Sulawesi, and Cluster 3 encompasses populations from Java, West Nusa Tenggara, and East Nusa Tenggara. These findings demonstrate high haplotype diversity and low nucleotide diversity within the species. Populations from West Sumatra, Manado, Tojo Una - Una, and North Morowali (Sulawesi) have the potential for speciation with a genetic distance of 0.61 – 0.94% for COI, between 0.81 – 0.95% for ITS2, and between 0.62 – 0.71% for COII. These findings underscore the need for further integrative studies to obtain a more comprehensive understanding of the *An. tessellatus* complex in Indonesia and its role in malaria transmission.

## 1. Introduction

*Anopheles tessellatus* (Theobald,1901) is recognized as a potentially significant vector of malaria and lymphatic filariasis across South, East, and Southeast Asia [1]. This taxon has been identified as the primary vector of pathogens responsible for these diseases in regions such as the Maldives, Sri Lanka, and certain parts of Indonesia. Taxonomically, the Tessellatus Group belongs to the Neomyzomyia Series of the *Anopheles* subgenus *Cellia.* It includes two species: *An. orientalis* and *An. tessellatus*, as well as the Tessellatus complex with six unnamed species A, B, C, D, E, and F [2, 3]. Originally described from specimens collected in Taiping, Perak, in Northwest Peninsular Malaysia [4], this species exhibits considerable complexity throughout Southeast Asia [2]. The taxonomic history of *An. tessellatus* is notably intricate, characterized by marked morphological diversity among populations, leading to the recognition of several synonyms and subspecies, particularly in Indonesia [5].

Morphological analyses have identified at least 11 morphological variants of *An. tessellatus* within the Indonesian archipelago. Among these, three subspecies have been formally recognized: *An. tessellatus tessellatus* (Theobald,1901); *An. tessellatus kalawara* (Stoker & Koesoemawinangoen, 1949), both described from Indonesia, and *An. tessellatus orientalis* (Swellengrebel and de Graaf, 1920), which is distributed across Sulawesi, Java, and the Maluku Islands [5]. The observed morphological diversity suggests the possible presence of cryptic lineages within the Tessellatus Complex, as recent molecular studies using COI gene analyses have identified up to six putative species within what was previously considered a single taxon [2]. Such cryptic diversity has significant implications for disease transmission dynamics and vector management, as divergent lineages may differ in their vectorial capacity and ecological characteristics [2].

Despite its epidemiological relevance and taxonomic complexity, studies on the genetic variation and population structure of *An. tessellatus* in Indonesia remains limited. Traditional morphological identification is often insufficient due to the isomorphic nature of sibling species, making molecular markers essential for accurate species delimitation and population genetic studies [6]. Among these, the Internal Transcribed Spacer 2 (ITS2) region of ribosomal DNA and the mitochondrial cytochrome c oxidase subunit I (COI) and subunit II (COII) genes have proven to be reliable tools for distinguishing closely related *Anopheles* species and assessing their genetic diversity [7]. Both ITS2 and COI markers have been widely used in population genetics, phylogenetic, and phylogeographic studies of *Anopheles* mosquitoes across Asia, revealing significant intra- and interspecific variation and uncovering cryptic species complexes [8]. Various molecular studies of *Anopheles* mosquitoes have also used the COI gene, including to identify members of the species complex. One example is the identification of the *An. subpictus* complex in Sri Lanka [9]. Studies on the genetic variation and population structure of *An. subpictus, An. peditaeniatus,* and *An. vagus* mosquitoes from five regions in Sri Lanka were also studied using the COI marker gene [6]. The COI gene, in particular, has been a cornerstone in studies addressing population structure and systematics of *Anopheles* mosquitoes [10, 11].

Previous studies have demonstrated the effectiveness of COI and ITS2 markers in identifying both Culicinae and *Anopheles* mosquitoes in diverse geographic settings, including Sri Lanka [6]. The ITS2 marker has been employed to elucidate species diversity and relationships among *Anopheles* populations in Malaysia, along the Laos-Cambodia border [12], where novel species within the *An. hyrcanus* group [13] and *An. lesteri in Korea* were discovered [14]. Similarly, investigations of *An. anthropophagus* populations in China have relied on ITS2 for species confirmation [12, 14] and a novel ITS2-based PCR was recently developed to better identify members of the *An. dirus* complex [15]. In addition to ITS2 and COI, the mitochondrial cytochrome c oxidase subunit II (COII) gene has emerged as a promising molecular marker for species identification and population genetic studies. For instance, studies on the *An. culicifacies* complex have successfully applied ITS2 and COII markers to resolve species boundaries [16]. Moreover, combined analyses of COI and COII genes have provided insights into the genetic population structure of *An. superpictus* in Iran [17].

Given these considerations, the present study aims to investigate the interpopulation genetic variation and population structure of *An. tessellatus* across Indonesia using molecular data from three genetic markers: the nuclear ITS2 ribosomal gene and the mitochondrial COI and COII genes. Specimens were collected from six geographically distinct populations representing Sumatra, Java, Kalimantan, the Lesser Sunda Islands, Sulawesi, and Papua. This thorough molecular approach is expected to provide a more comprehensive understanding of the genetic diversity, evolutionary relationships, and potential vectorial capacity of *An. tessellatus* in Indonesia, thereby contributing valuable information for malaria and lymphatic filariasis control programs in Indonesia/the studied area.

## 2. Methods

### Sample collections and identification

Adult mosquitoes were systematically collected from diverse field environments across 13 Indonesian provinces between 2015 and 2024, employing standardized human landing catches, cattle-bait collections, US CDC light traps, and animal-baited trap techniques. Sampling sites encompassed the South Coast (Pesisir Selatan) in West Sumatra; Central Sumba and Southwest Sumba in East Nusa Tenggara; West Lombok and Bima in West Nusa Tenggara; Manado in North Sulawesi; Tojo Una-Una, Banggai, North Morowali, Sigi, Palu, and Donggala in Central Sulawesi; West Coast (Pesisir Barat) in Lampung; Murung Raya in Central Kalimantan; and Kulon Progo in the Special Region of Yogyakarta. (See Fig. 1; Table 1). Initial identification of *An. tessellatus* specimens were performed based on morphological characteristics. Collected mosquitoes were sorted and labelled according to location and collection date, then preserved in 1.5 ml Eppendorf tubes containing silica gel to maintain dry conditions until further analysis [18]. Samples were obtained from field mosquito collections and access to stored biological materials (*Anopheles tessellatus* mosquitoes) was obtained from the Center for Environmental Health Laboratory, The Ministry of Health, Indonesia. Permissions to access these samples for research purposes were secured via a Material Transfer Agreement authorized by the Director of the Center for Environmental Health Laboratory (no. HK.03.01/IX.1/3787/2024) and the Dean of the Faculty of Biology, Universitas Gadjah Mada (no. 6941/UN1/FBI/KSA/HK.08.00/2024) on December 14, 2024.

**Figure 1.**
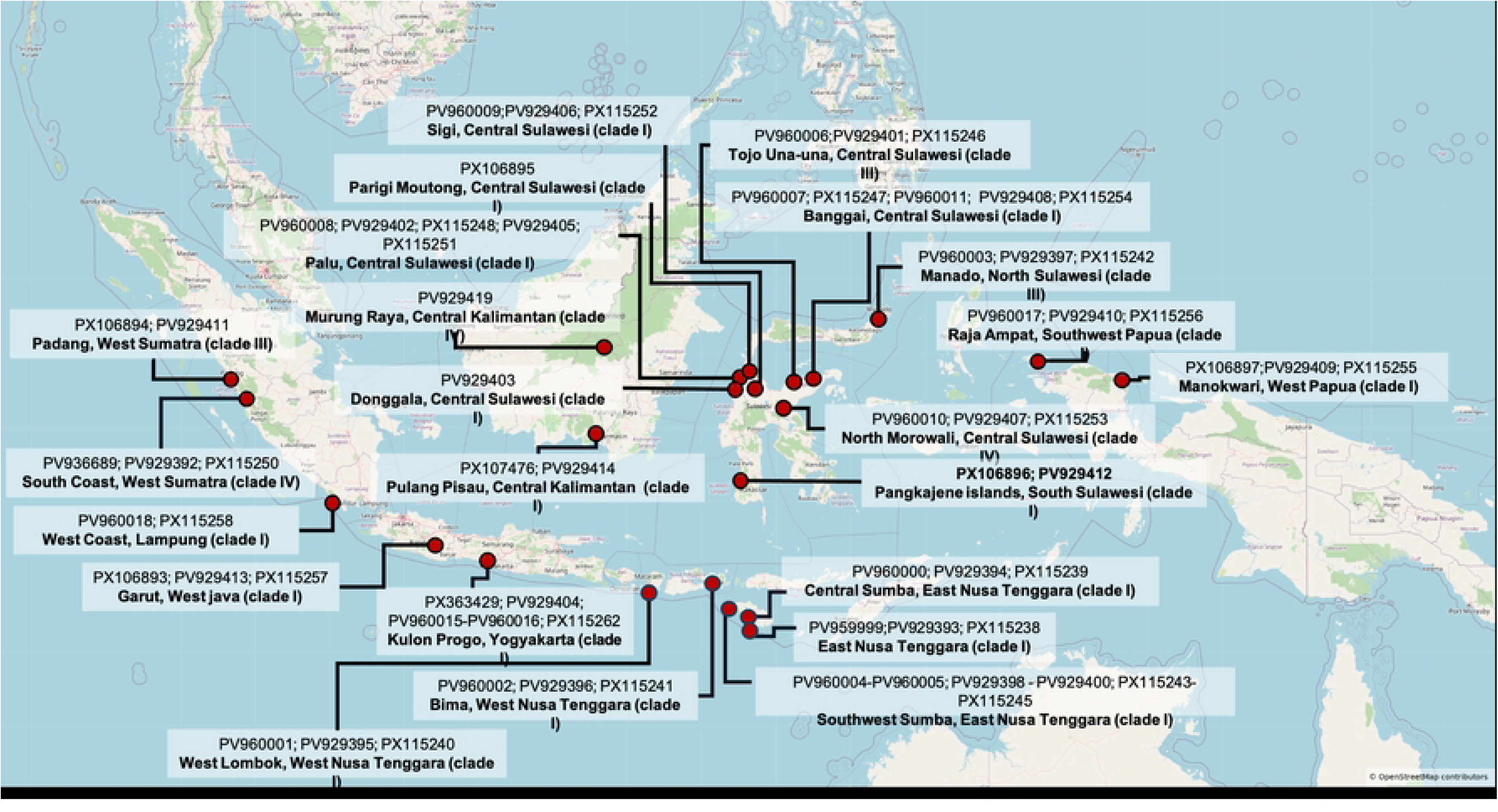
Distribution map of *Anopheles tessellatus* sampling locations in Indonesia, marked with red dots.

**Table 1.**
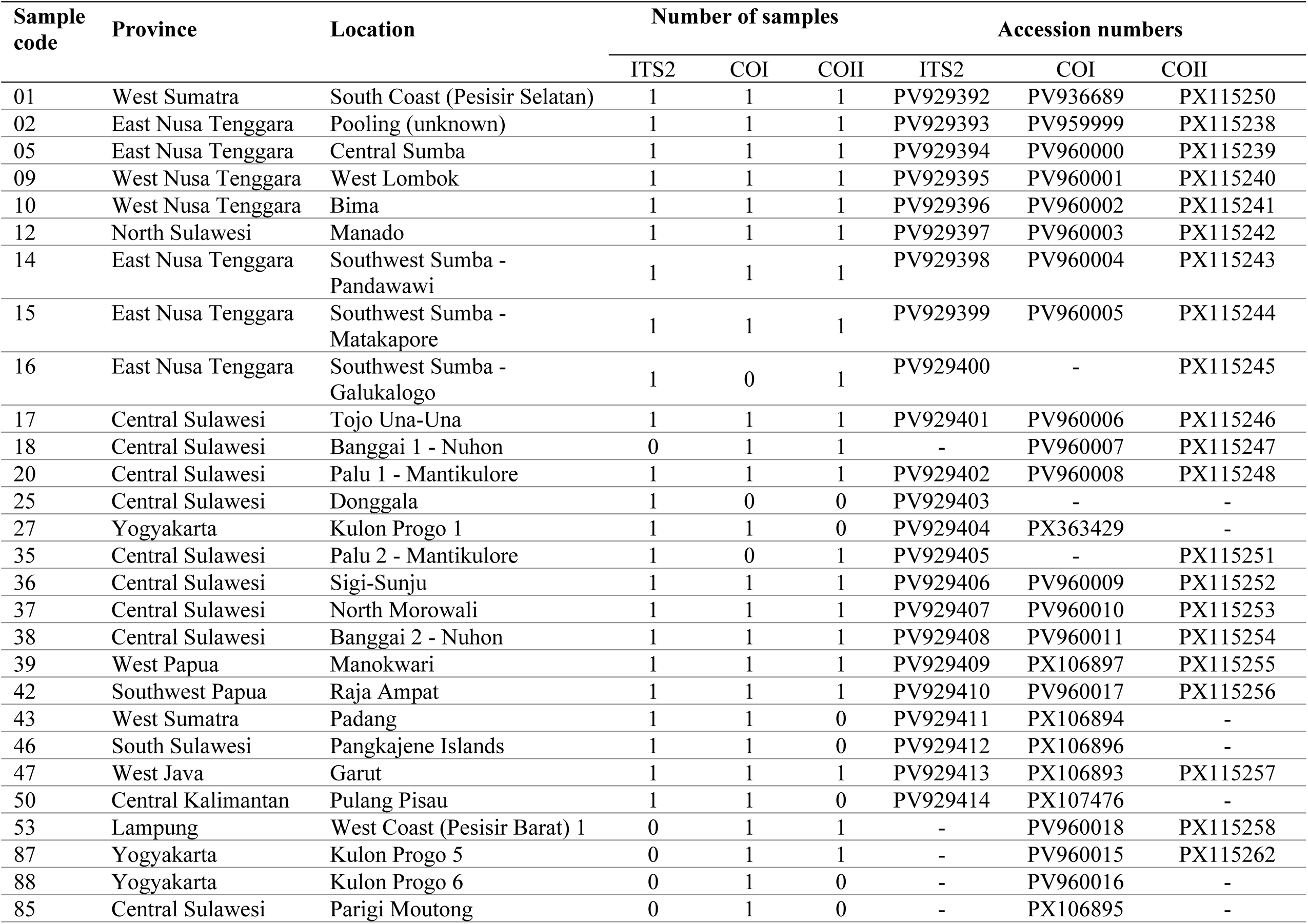
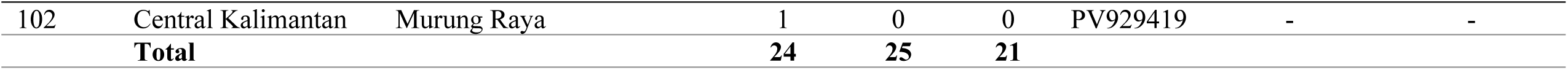
Sampling localities and specimens of the *Anopheles tessellatus* complex (Number sequenced)

### DNA extraction, amplification, and sequencing

Genomic DNA was isolated from individual mosquito samples using the Zymo Quick-DNA Miniprep Plus DNA extraction kit (ZYMO, Irvine, CA, USA) according to the manufacturer’s protocols. Three genetic loci were targeted for PCR amplification: the internal transcribed spacer 2 (ITS2) region, cytochrome c oxidase subunit I (COI), and cytochrome c oxidase subunit II (COII). ITS2 was amplified with primers ITS2a (5’-TGTGAACTGCAGGACACAT-3’) and ITS2b (5-TATGCTTAAATTCAGGGGGT-3’), COI with primers LCO1490 (5’-GGTCAACAAATCATAAAGATATTGG-3’) and HCO2198 (5’-AAACTTCAGGGTGACCAAAAAATCA-3’), and COII with primers Anplcox2F (5’-GGATCCAGATTAGTGCAATGAATTTAAGC-3’) and Anplcox2R (5’-CTGCAGGATTTAAGAGATCATACTTGC-3’). PCR reactions were performed using GoTaq Green Master Mix (Promega, Madison, WI, USA). Thermocycling conditions for ITS2 consisted of an initial denaturation at 94 °C for 10 min, followed by 40 cycles of 94 °C for 1 min, 52 °C for 45 s, and 72 °C for 1 min, with a final extension at 72 °C for 10 min. For COI, the protocol included an initial denaturation at 94 °C for 1 min; five cycles of 94 °C for 30 s, 45 °C for 40 s, and 72 °C for 1 min; followed by 35 cycles of 94 °C for 30 s, 55 °C for 40 s, and 72 °C for 1 min; and a final extension at 72 °C for 10 min. COII amplification followed the same cycling parameters as COI, except the annealing temperature during the 35 cycles was set at 58 °C. PCR products were separated by electrophoresis on 1.5% agarose gels, visualized using SYBR Safe DNA gel stain (Invitrogen, Carlsbad, CA, USA), and fragment sizes were estimated with a 100 bp DNA ladder. PCR amplification products were then purified using the ExoSAP-IT™ reagent (Applied Biosystems, Thermo Fisher Scientific, Vilnius, Lithuania) prior to sequencing. Sequencing reactions were performed using the same primers described above, with the BigDye^TM^ Terminator v3.1 Cycle Sequencing Kit (Life Technologies Corporation, Austin, TX, USA). Resulting sequences were processed and edited using Sequencing Analysis software version 5.2 (Applied Biosystems). Multiple sequence alignments were generated with ClustalW 1.6 implemented in MEGA X version 10.2.2. The obtained sequences were compared against reference sequences of *Anopheles tessellatus* available in the NCBI GenBank database. Multiple sequence alignment was performed using MUSCLE v.5.3 [19]. The aligned sequences were further trimmed with trimAl v.1.5 [20] using the “-automated” setting, which was optimized for Maximum Likelihood (ML) phylogenetic tree reconstruction. ML trees were inferred using IQ-TREE v.2.3.6 [21] with optimal models selected from the full range of models supported by ModelFinder [22]. Clade support was assessed with 1,000 bootstrap replicates using UFBoot2 [23]. Hierarchical cluster analysis was performed to identify clusters in the reconstructed phylogenetic trees using R v.4.2.2 [24]. Pairwise p-distance distances were calculated using the APE R package [25]. The minimum interspecific and maximum intraspecific distances were calculated using the SPIDER R package [26]. The standard genetic indices, haplotype diversity (h), haplotype number (H), and nucleotide diversity (π) were evaluated using DnaSP v.6.12.03 [27]. Haplotype networks were constructed with PopArt [28] using the TCS estimating gene genealogy algorithm [29].

### Ethical Considerations

This study involved human participants who collected adult mosquitoes in natural field settings. Formal approval for these activities was granted by the Health Ethics Commission of the National Research and Innovation Agency (BRIN), under approval number 111/ KE.03/SK/05/2024.

## 3. Results

### The Internal Transcribed Spacer 2 (ITS2) diversity, phylogeny, and polymorphism of *Anopheles tessellatus*

Analysis of ITS2 rDNA sequences from 24 *An. tessellatus* samples collected at 23 locations across Indonesia, supplemented by 30 reference sequences from GenBank (23 sequences from Thailand, three sequences from Indonesia, three from Malaysia, and one from China), revealed the presence of four distinct *An. tessellatus* populations within Indonesia (Table 1, Fig. 1, Fig. 2). The majority of study samples clustered within Clade I, encompassing specimens from Kulon Progo (Yogyakarta), Pulang Pisau (Central Kalimantan), Garut (West Java), Pangkajene Islands (South Sulawesi), Banggai, Sigi, Donggala, and Palu (Central Sulawesi), West Lombok and Bima (West Nusa Tenggara), Galukalogo, Matakapore, Pandawawi, and Central Sumba (East Nusa Tenggara), Raja Ampat (Southwest Papua), Waropen, and Manokwari (West Papua) (Accession numbers: PV929394-PV929396, PV929398-PV929400, PV929402-PV929406, PV929408-PV929410, PV929412-PV929414). Genetic distances within Clade I ranged from 0% to 0.13%, with the greatest divergence observed between the Garut sample and sequence from Clade II (the genetic distance within Clade II range from 0% to 0.95%), a clade predominantly composed of samples from Thailand and one from China, with no Indonesian samples (Fig. 2; Fig. 3).

**Figure 2.**
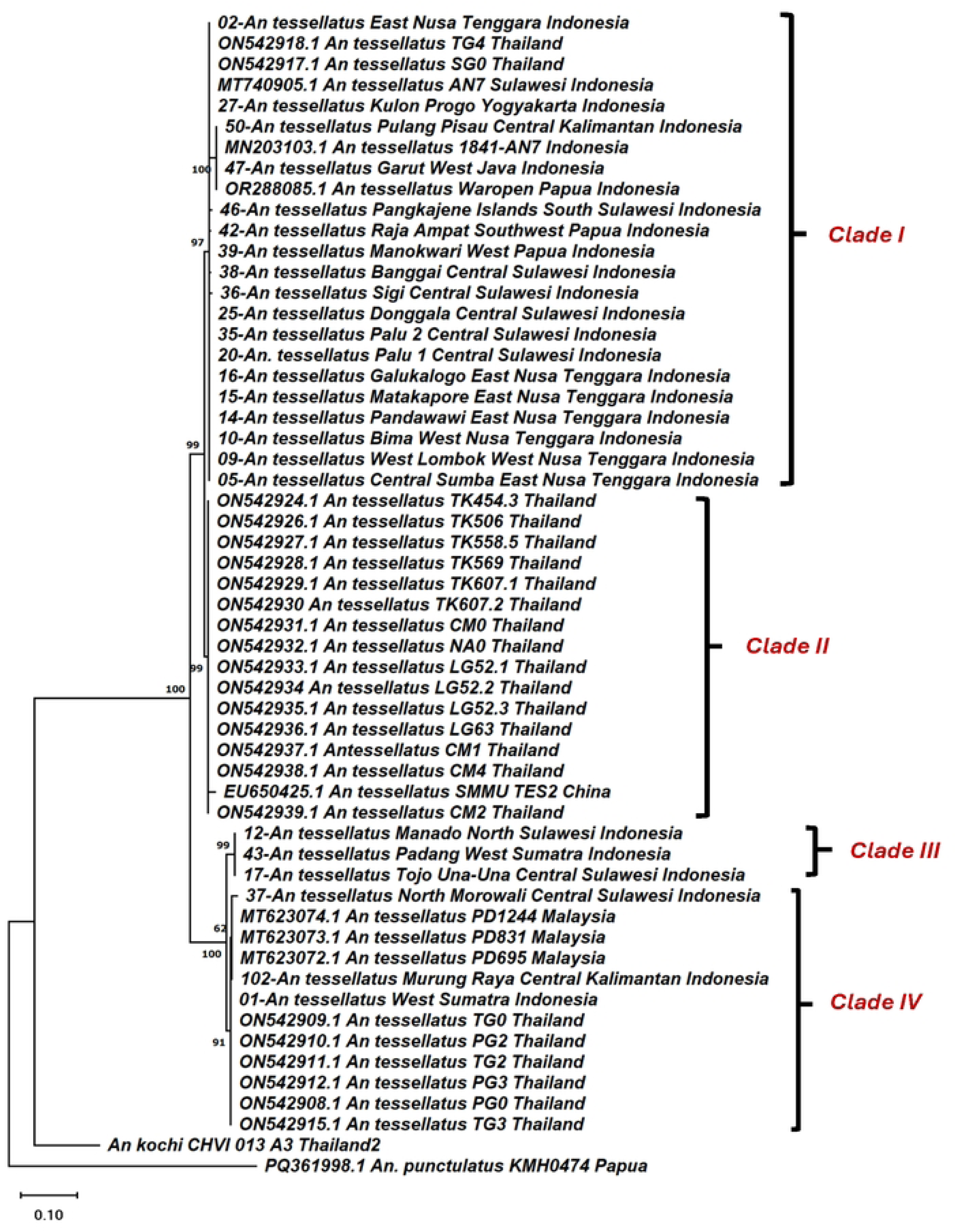
Phylogenetic analysis of the ITS2 sequences of the *Anopheles tessellatus* complex, rooted with *Anopheles punctulatus*. The phylogenetic tree was constructed using the Maximum Likelihood (ML) method with the Kimura 2-parameter evolutionary model in MEGA X, and bootstrap values were tested with 1,000 replicates.

**Figure 3.**
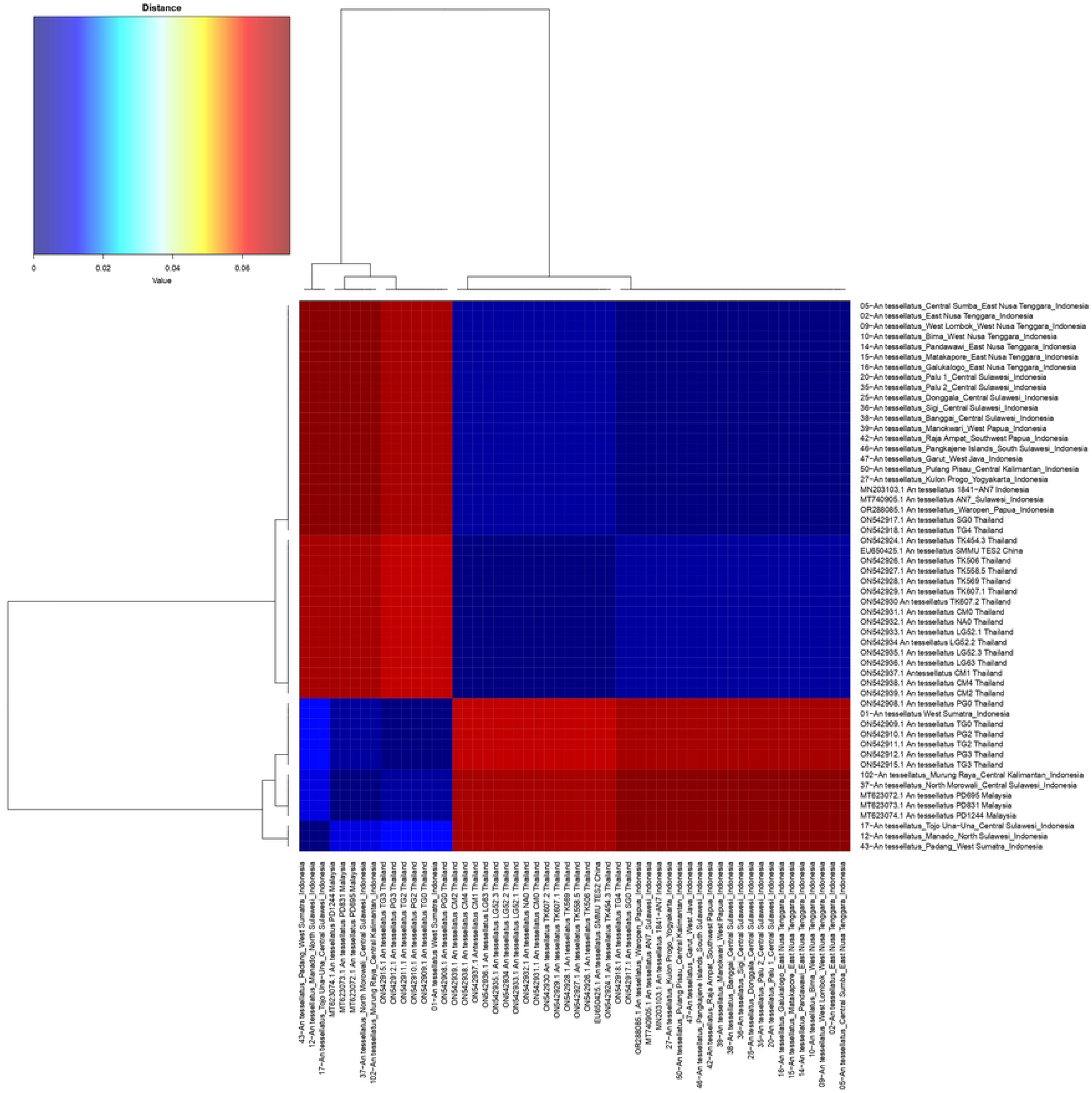
Pairwise nucleotide genetic distances (p-distance model) of *Anopheles tessellatus* ITS2 sequences.

Clade III, notably distinct, comprised samples from geographically distant locations: Padang (West Sumatra), Tojo Una-Una (Central Sulawesi), and Manado (North Sulawesi) (PV929396, PV929401, PV929411). Intra-clade genetic distances ranged from 0.14% to 1.14%, with the Padang samples exhibiting the highest divergence. Clade IV included samples from the South Coast of West Sumatra, North Morowali (Central Sulawesi), and Murung Raya (Central Kalimantan) (Accession numbers: PV929392, PV929407, PV929414), with genetic distances within this group ranging between 0.05% and 0.09% (Fig. 2; Fig. 3: Table 2).

Despite the presence of four genetically distinct populations, inter-clade genetic distances ranged from 0.01% to 1.03%. Clade I demonstrated the greatest diversity and the broadest geographic distribution across Indonesia. Clade II appears to represent a population widely distributed in Thailand and China. Samples assigned to Clade III correspond to the Sulawesi lineage, currently identified only on the island of Sulawesi. Clades I and III represent unique populations endemic to Indonesia. Conversely, Clade IV displays a wider distribution across three Southeast Asian countries, including Malaysia, Indonesia, and Thailand (Fig. 2; Fig. 3; Table 2).

ITS2 haplotype analysis reveals distinct genetic diversity patterns in *An. tessellatus* populations across Indonesia and reference sequences from other regions in Asia (Fig. 4). Five distinct haplotypes were identified from 54 sequences, supported by two variable sites, with a haplotype diversity (Hd) of 0.716, indicating moderate intraspecific variation suitable for population studies in vector mosquitoes [30]. Hap1, the most frequent (n=7), includes sequences from West Sumatra (Indonesia) and Thailand, differing at positions 32-37, suggesting a widespread ancestral lineage. Hap2 (n=23) dominates Indonesian samples from East Nusa Tenggara, West Nusa Tenggara, Central Sulawesi, West Papua, South Sulawesi, West Java, Central Kalimantan, and Yogyakarta, with mutations at positions 2-5, 7-9, 11-14, 16-18, 20-23, 26, 30-31, and 38-39, reflecting regional diversification. Hap3 (n=3) appears limited to North and Central Sulawesi (Manado, Tojo Una-Una), and West Sumatra, varying at sites 6, 10, and 19. Hap4 (n=5) links Central Sulawesi, Central Kalimantan, and Malaysia (positions 15, 24, 27-29). Hap5 (n=16), primarily Thai sequences with one Chinese reference, shows extensive variation at sites 25 and 40-54, highlighting geographic structuring. Haplotype networks, typically constructed via median-joining or TCS methods, visualize mutational steps between haplotypes, with Hap1 as a central node linking Indonesian-Thai populations and radiating to Hap2 (most diverged, multi-site changes) and minor haplotypes (Hap3-5). The presence of short mutational branches among Indonesian Hap2 sequences indicate recent population expansion or gene flow across islands, while longer branches to Hap5 suggest historical isolation or concerted evolution minimizing ITS2 intraspecific polymorphism. This structure implies that *An. tessellatus* comprises cryptic lineages, with Hap2 potentially adapted to diverse Indonesian ecologies, warranting vector competence assessments for malaria control.

**Figure 4.**
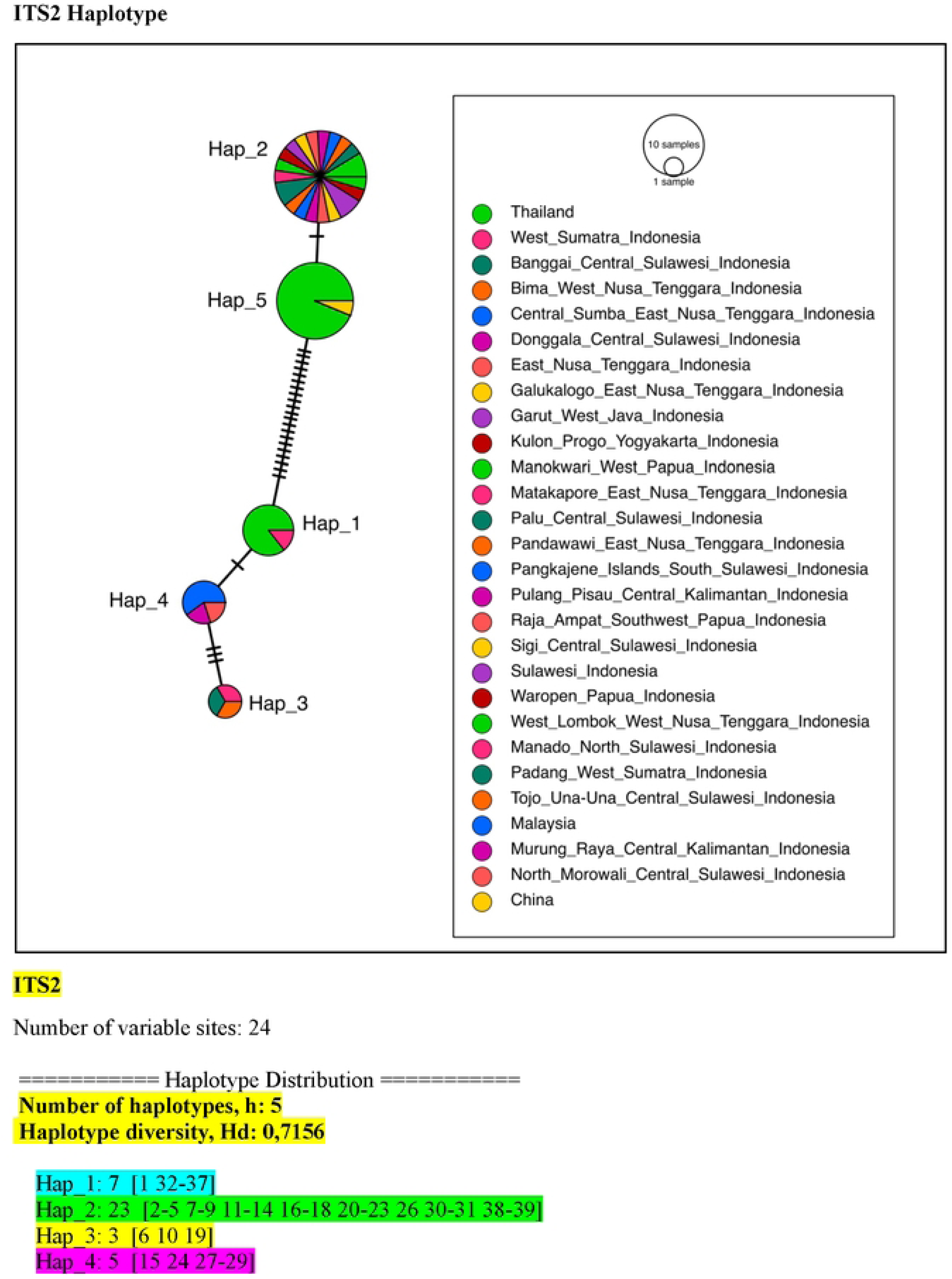
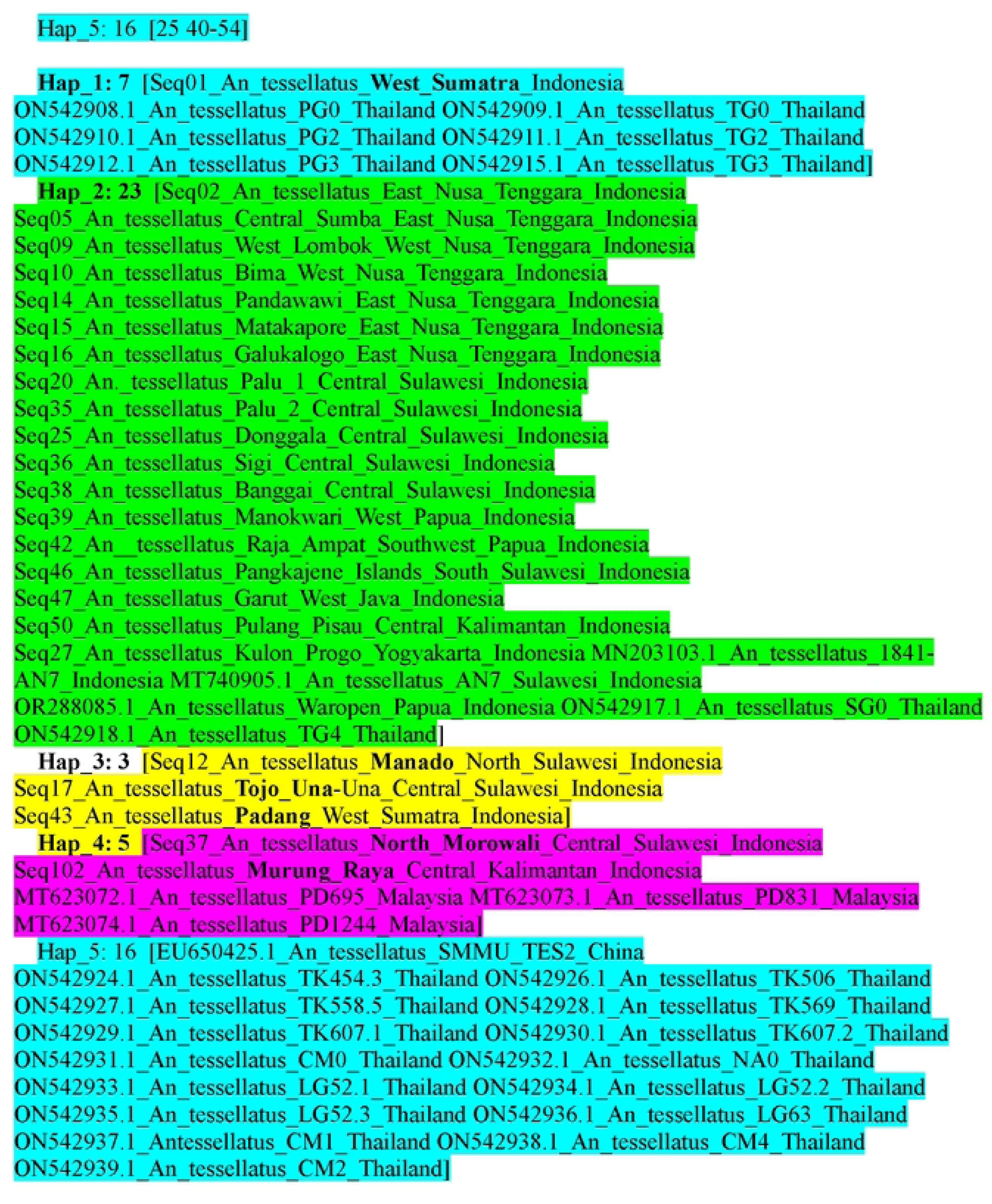
Haplotype network of *Anopheles tessellatus* based on ITS2 sequences. The figure illustrates the frequency of each ITS2 haplotype across study sites, with connections representing the number of mutations separating haplotypes and their geographic distribution.

### The Cytochrome Oxidase I gene (COI) diversity, phylogeny, and polymorphism of *Anopheles tessellatus*

Analysis of COI sequences to identify maternal lineages revealed that all 25 samples from this study (22 locations across Indonesia), together with 31 reference sequences from GenBank (five from Indonesia, three from Malaysia, four from Singapore, five from Philippines, five from Thailand, three from Sri Lanka, two from Japan, three from Vietnam, one from Laos) grouped into 12 distinct lineages within the *An. tessellatus* populations. Lineage 1 comprised samples from Kulon Progo (Yogyakarta) (PX 363429), Banggai, Sigi, and Palu (Central Sulawesi (PV960007, PV960009, PV960008, PV960011), Pandawawi, Central Sumba, and Matakapore (East Nusa Tenggara) (PV960004, PV960000, PV960005), Bima and West Lombok (West Nusa Tenggara) (PV960002, PV960001), Raja Ampat and Manokwari (West Papua) (PV960017, PX106897), along with six samples of GenBank reference sequences from East Kalimantan (MT257012), Sulawesi (MT753038), Sarawak (MT256986, MT256985, MT256984), and the Philippines (MT256990) (Table 1, Fig. 5).

**Figure 5.**
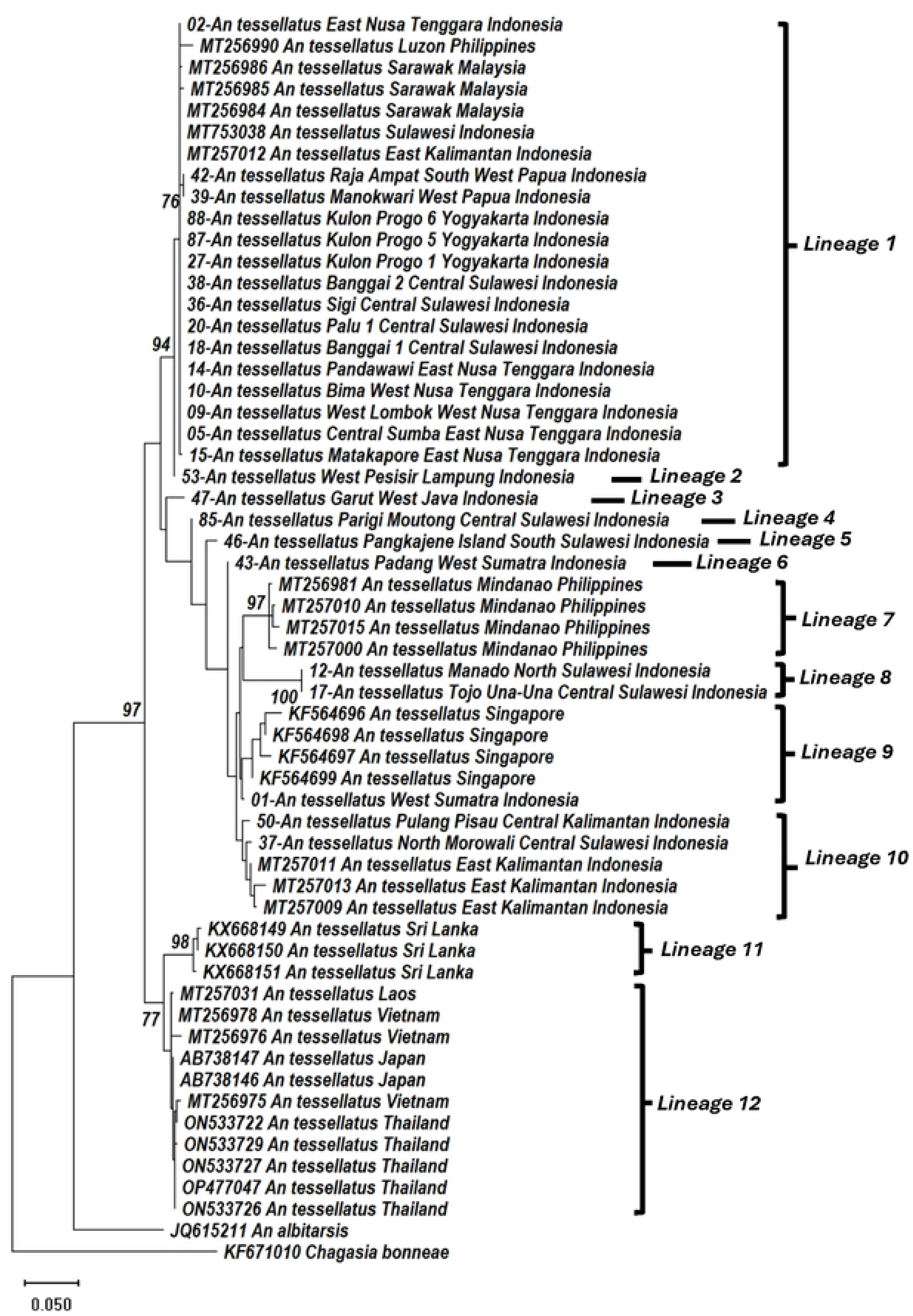
Phylogenetic analysis of COI sequences of *Anopheles tessellatus*, rooted with *Chagasia bonneae* as the outgroup. The phylogenetic tree was constructed using the General Time Reversible (GTR+I) model in MEGA X, with bootstrap support estimated from 1,000 replicates to assess tree reliability.

Lineage 2 was represented solely by specimens from West Pesisir, Lampung (PV960018), while lineage 3 contained only sample from Garut (West Java) (PX106893). Subsequently, lineages 4, 5, and 6, each contained samples from Parigi Moutong (Central Sulawesi) (PX106895), Pangkajene Islands (South Sulawesi) (PX106896), and Padang (West Sumatra) (PX106894), respectively. Lineage 7 consisted exclusively of GenBank references from the Philippines (MT256981, MT257000, MT257010, MT257015). Lineage 8 included specimens from Manado (North Sulawesi) (PV960003) and Tojo Una-Una (Central Sulawesi) (PV960006). Lineage 9 encompassed samples from the South Coast of West Sumatra (PV936689) and four GenBank references from Singapore (KF564696-KF564699). Lineage 10 contained specimens solely from Indonesia, including Pulang Pisau (Central Kalimantan) (PX107476), GenBank references from East Kalimantan (MT257009, MT257011, MT257013), and North Morowali (Central Sulawesi) (PV960010). Lineage 11 was represented only by three GenBank samples from Sri Lanka (KX668149-KX668151), whereas lineage 12 comprised a mixture of GenBank sequences from four Asian countries: Thailand (ON533722, ON533726, ON533727, ON533729, OP477047), Japan (AB738146-AB738147), Vietnam (MT256975-MT256976, MT256978), and Laos (MT257031). Phylogenetic rooting was achieved using *Chagasia bonneae* (KF671010) and *Anopheles albitarsis* (JQ615211) as outgroups (Fig. 5).

The majority of *An. tessellatus* samples in this study fell within Lineage 1, exhibiting minimal genetic divergence with a distance ranging from 0% to 0.04%. Apart from Lineage 1, Indonesian specimens also clustered in lineages 2, 3, 4, 5, 6, 8, 9, and 10. Notably, samples had genetic distances ranging from 0.04% to 0.6%. Inter-lineage COI sequence divergences ranged between 0.03% and 0.6%. These findings indicate that despite geographic separation across various regions and islands in Indonesia over extended evolutionary timescales, specifically since at least the late Jurassic Xenozoic era, approximately 2 million years ago[31]. *An. tessellatus* populations in Indonesia remain part of a single maternal lineage (Fig. 5, Fig. 6; Table 3).

**Figure 6.**
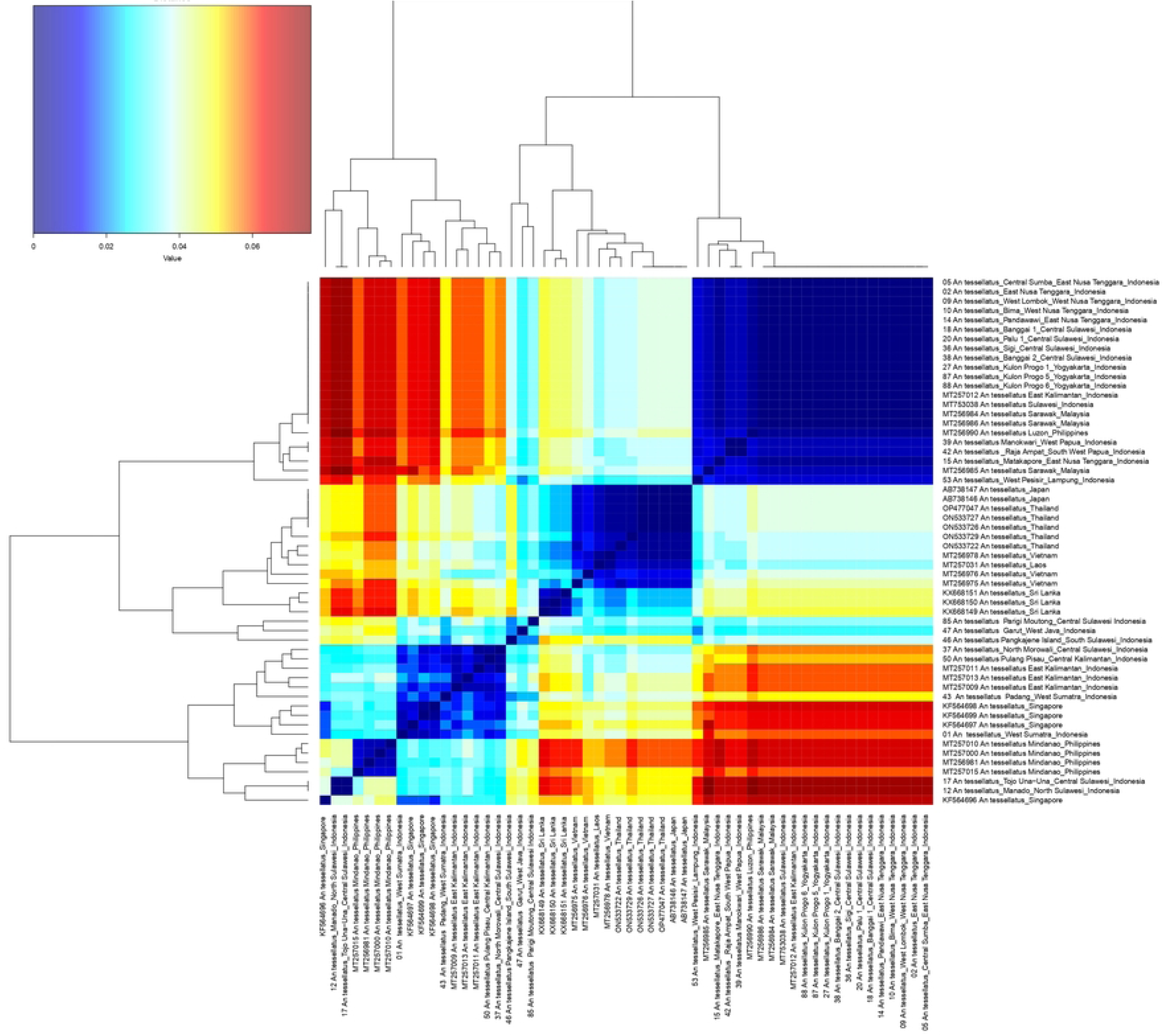
Pairwise nucleotide genetic distances (p-distance model) of *Anopheles tessellatus* COI sequences.

Subsequently, COI haplotype analysis of *An. tessellatus* demonstrates exceptionally high genetic diversity, with 35 haplotypes identified from approximately 54 sequences across 56 variable sites, resulting a haplotype diversity (Hd) of 0.914 (Fig. 7). This high level of haplotype diversity underscores the sensitivity of the mitochondrial COI marker to recent demographic processes in mosquito populations and exceeds the diversity detected using ITS2, thereby reaffirming the utility of COI for detecting cryptic genetic structure within *An. tessellatus* across Southeast Asia [2]. Hap2 was the most dominant haplotype (n=16), encompassing Indonesian samples from East Nusa Tenggara, West Nusa Tenggara, Central Sulawesi, Yogyakarta, as well as references from East Kalimantan (Indonesia) and Sarawak (Malaysia). This haplotype is defined by mutations at positions 2-5, 7, 10-12, 14-17, 28, 30-31, and 33, indicating a widespread, possibly ancestral clade with ongoing gene flow [6]. Hap27 (n=5) clusters Japanese and Thai references (positions 44-45, 52, 54-55), while singleton haplotypes (e.g., Hap1 from West Sumatra; Hap3 n=2 from North /Central Sulawesi at sites 6,9) predominate, alongside unique regional variants from Papua, West Java, Lampung, Kalimantan, Philippines, Singapore, Sri Lanka, Vietnam, and Laos, underscoring pan-Asian phylogeographic structuring. In median-joining or statistical parsimony networks, Hap2 is typically positioned as a central node, connected by short mutational branches to Indonesian singletons (Hap 4-6, 9-12), reflecting low divergence and substantial connectivity likely facilitated by island dispersal or anthropogenic movement. In contrast, longer branches leading to Hap27 (Thailand/Japan) and to more distantly related Asian haplotypes (e.g., Hap28-35), implying historical vicariance or isolation by distance. His star-like topology with a dominant hub suggests recent demographic expansion in Indonesian populations, contrasting lower ITS2 diversity and aligning with neutral evolution under Tajima’s D neutrality tests for vector surveillance [2].

**Figure 7.**
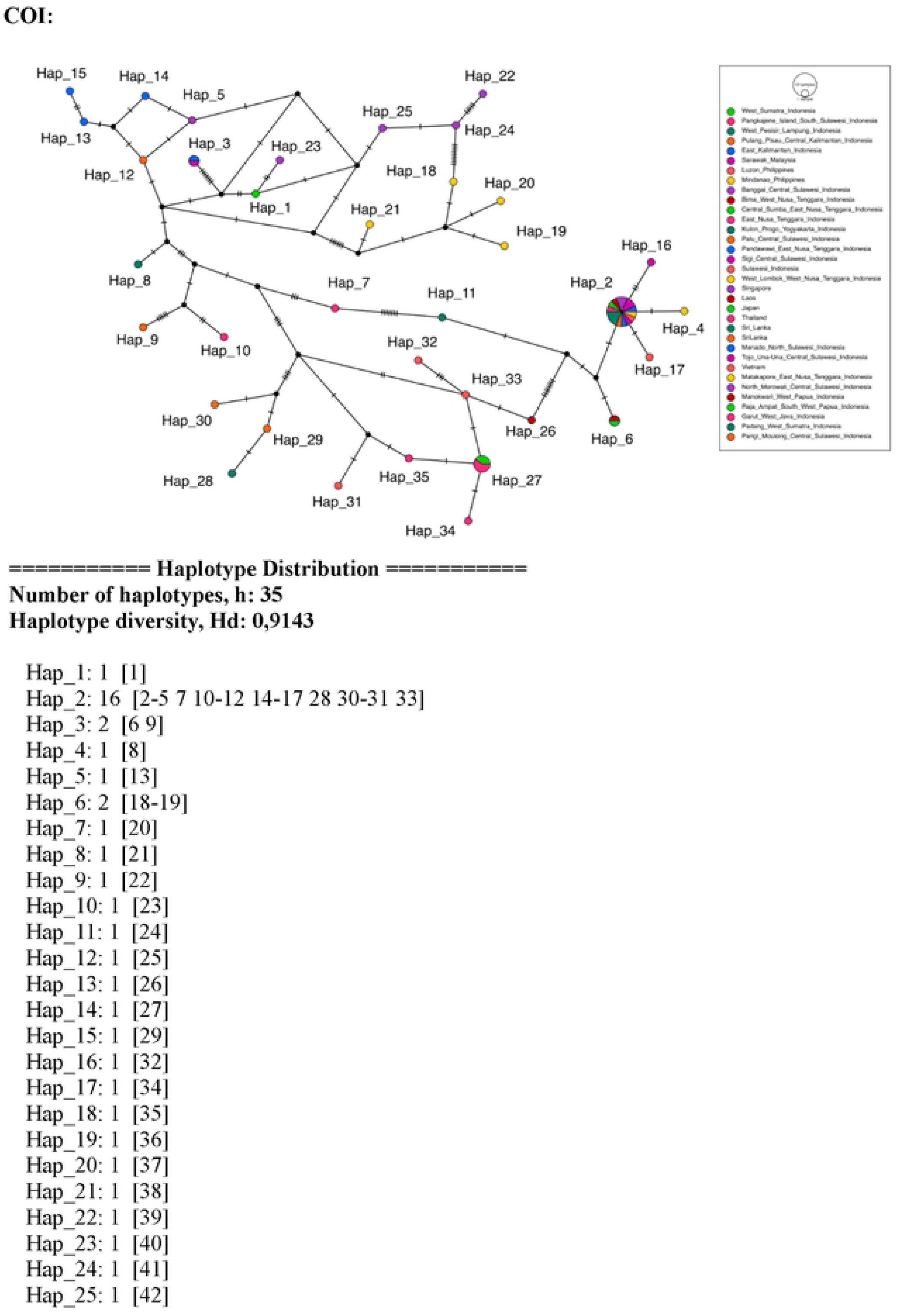
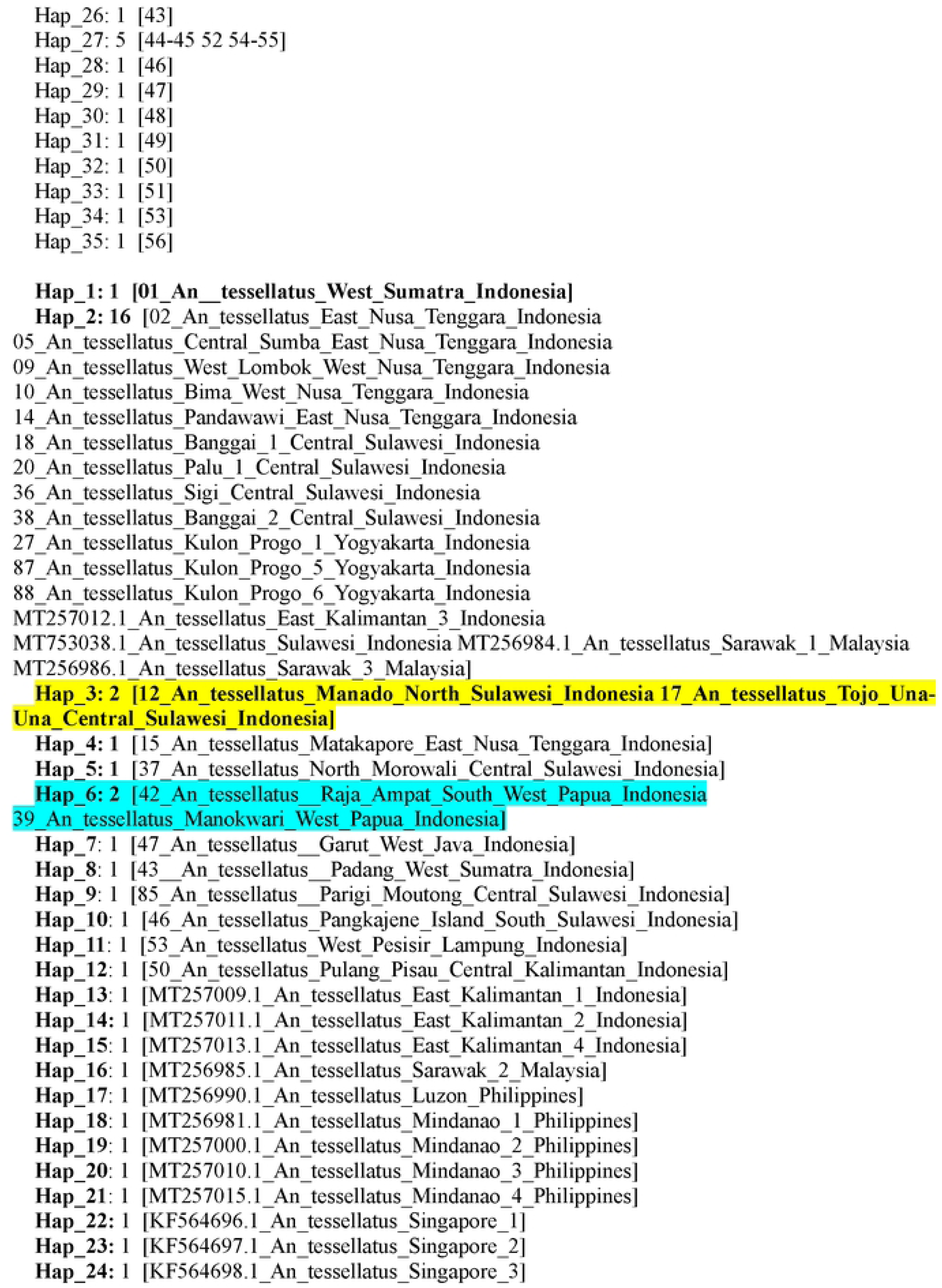
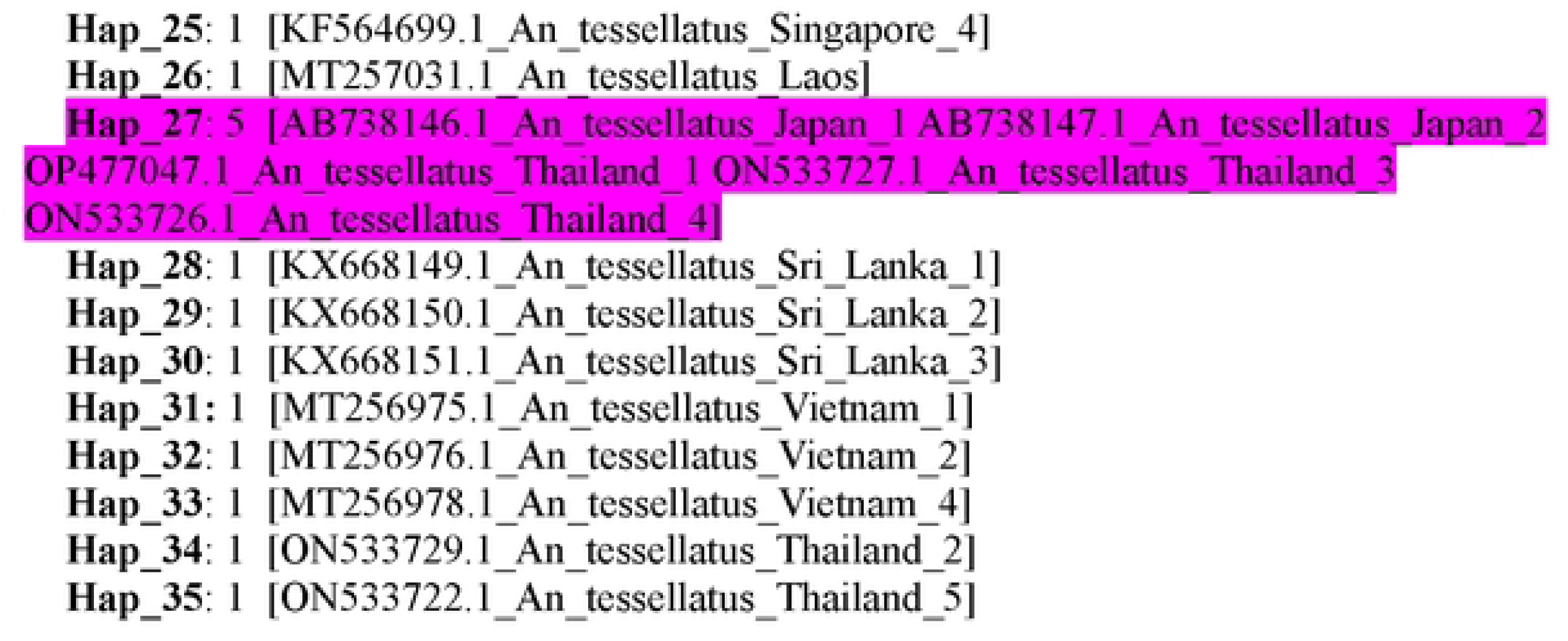
Haplotype network of *Anopheles tessellatus* based on COI sequences. The figure depicts the frequency of each COI haplotype across the study sites, with lines indicating the number of mutational steps separating haplotypes and their geographic distribution.

### The Cytochrome Oxidase II gene (COII) diversity, phylogeny, and polymorphism of *Anopheles tessellatus*

Analysis of COII sequences from 21 *An. tessellatus* specimens collected in this study (18 locations) across Indonesia, along with eight reference sequences retrieved from GenBank (one from Australia, two from China, and five from Thailand), revealed four distinct population clusters. Cluster I comprised specimens from West Pesisir (Lampung) (PX 115250), Sigi, Banggai, and Palu (Central Sulawesi) (PX115252, PX115247, PX115254, PX115248, PX115251), Garut (West Java) (PX115257), Kulon Progo (Yogyakarta) (PX115262), West Lombok and Bima (West Nusa Tenggara) (PX115240-PX115241), Pandawawi, Matakapore, Galukalogo, and Central Sumba (East Nusa Tenggara) (PX115243-PX115245, PX115239), Manokwari (PX115255), Raja Ampat (Southwest Papua) (PX115256), and one sample of GenBank reference from Australia (U94315). Cluster II consisted of GenBank reference samples from Thailand (ON455345-ON455349) and China (EU620674). Cluster III contained specimens from West Sumatra (PX115250), North Morowali (Central Sulawesi) (PX115253), Tojo Una-Una (Central Sulawesi) (PX115246), and Manado (North Sulawesi) (PX115242), Cluster IV included a single GenBank reference sequence from China (JX070727) (Table 1; Fig. 8; Table 4).

**Figure 8.**
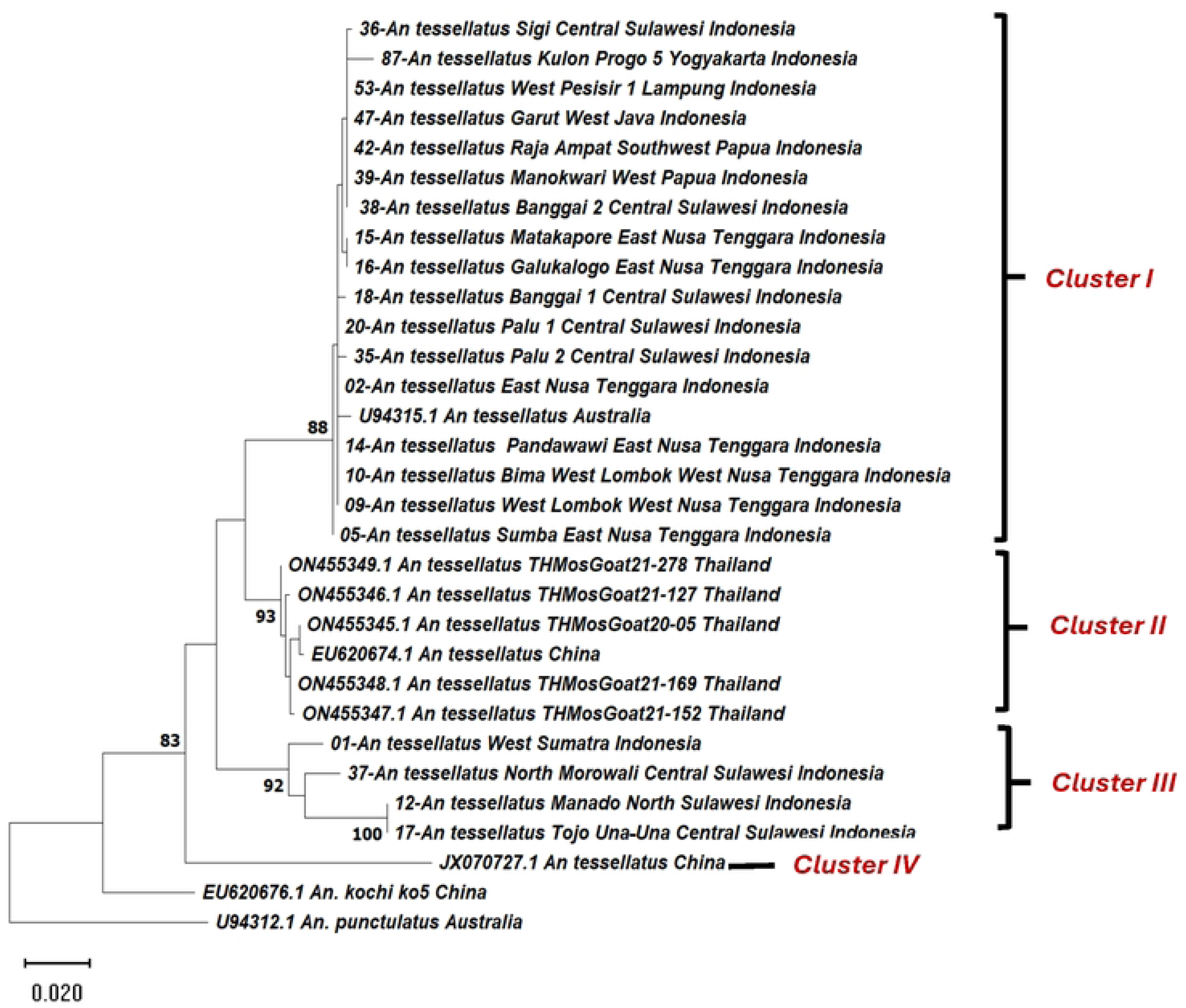
Phylogenetic analysis of COII sequences of *Anopheles tessellatus*, rooted with *Anopheles punctulatus* as the outgroup. The phylogenetic tree was constructed using the General Time Reversible (GTR+I) model in MEGA X, with bootstrap support estimated from 1,000 replicates to assess tree reliability.

Genetic distances among members of Cluster I ranged from 0.02% to 0.13%, whereas intra-cluster COII sequence divergence within Cluster II ranged from 0.01% to 0.06%. In Cluster III, genetic distances were higher, ranging from 0.24% to 0.33%. Overall COII divergence among all samples ranged from 0.03% to 0.36%. The COII phylogenetic analysis revealed a simpler clustering pattern that was nonetheless congruent with the COI-based phylogeny, indicating that *An. tessellatus* populations distributed across Indonesia remain within the same maternal lineage despite long-term geographic isolation among islands and regions. The greater apparent complexity of the COI analysis likely reflects the larger number and broader taxonomic and geographic representation of *An. tessellatus* references available in GenBank for COI compared with COII (Fig. 9; Table 4).

**Figure 9.**
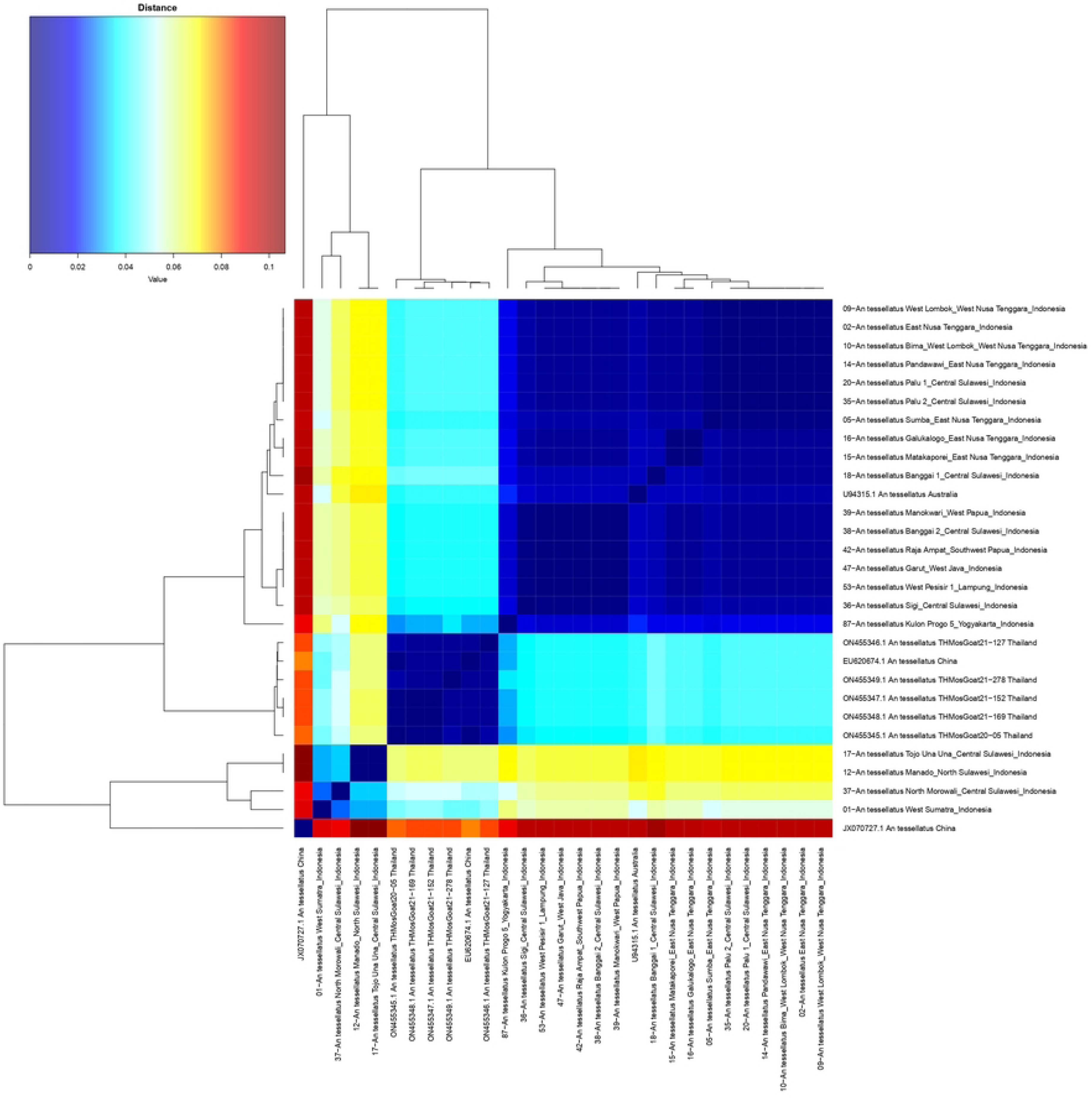
Pairwise nucleotide genetic distances (p-distance model) of *Anopheles tessellatus* COII sequences.

**Figure 10.**
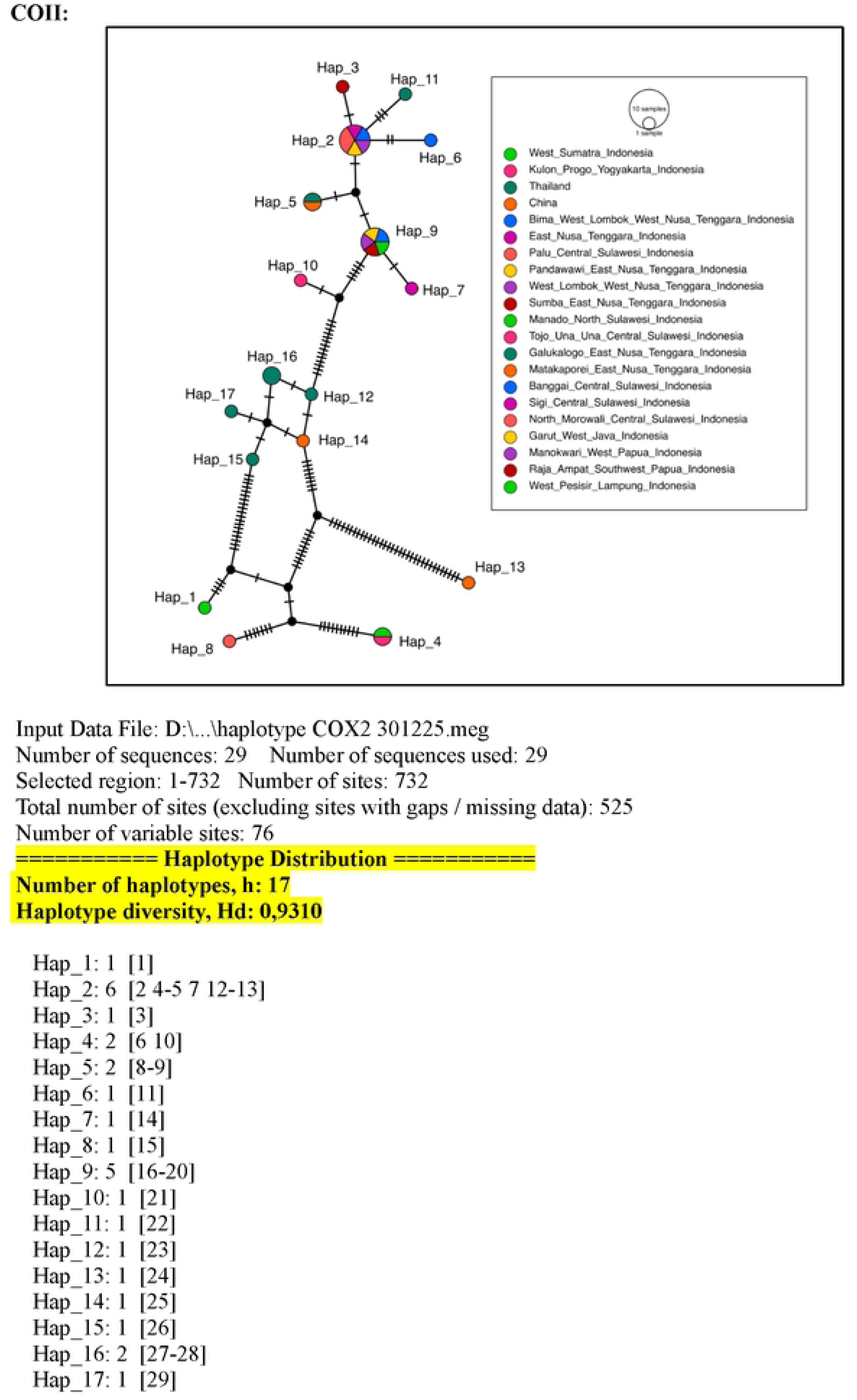
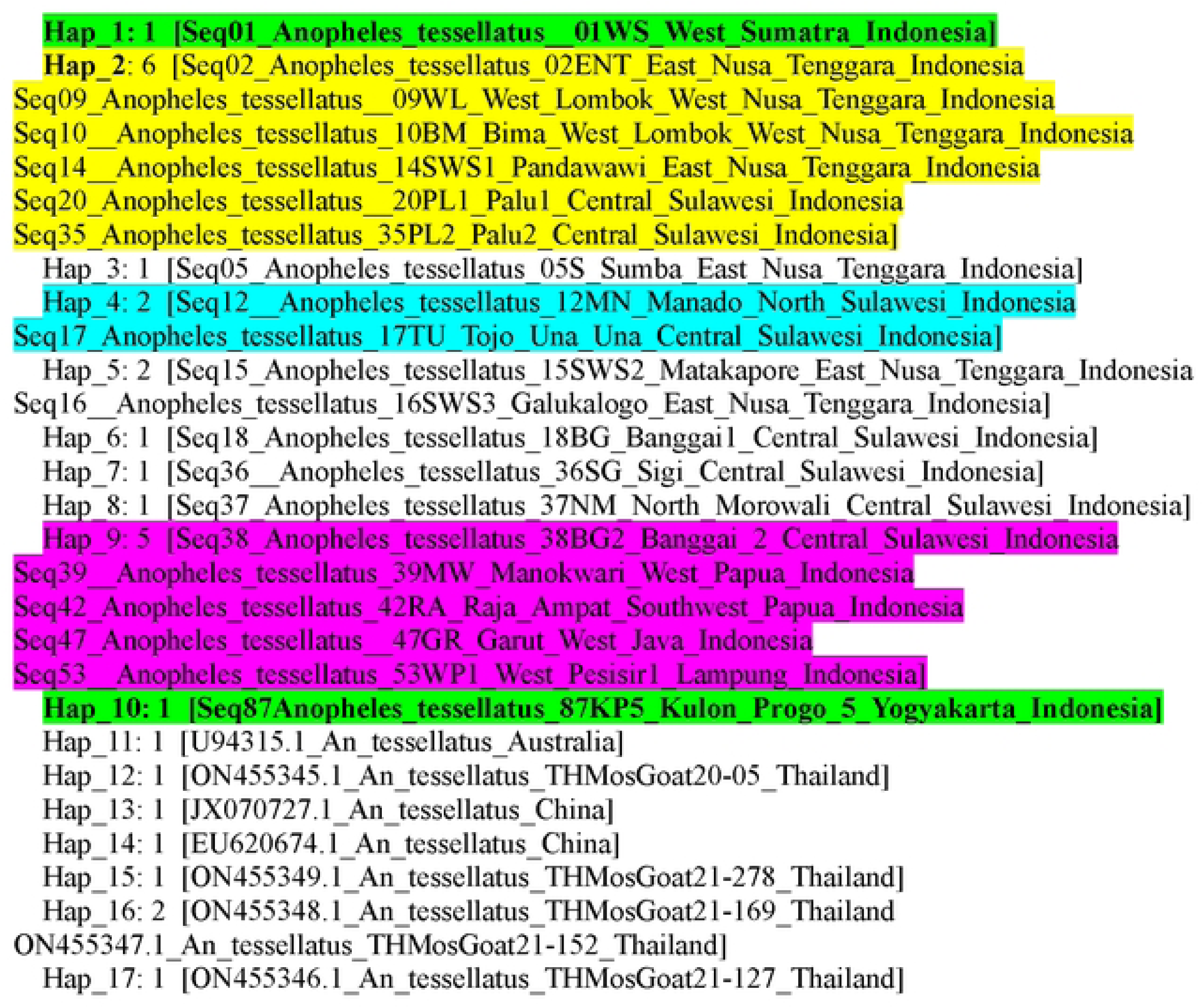
Haplotype network of *Anopheles tessellatus* based on COII sequences. The figure shows the frequency of each COII haplotype across the study sites, with connecting lines representing the number of mutational steps separating haplotypes and their geographic distribution.

Furthermore, COII haplotype analysis of *An. tessellatus* based on 29 Indonesian sequences (525 bp alignment, 76 variable sites) reveals high genetic diversity, with 17 haplotypes and haplotype diversity (Hd) of 0.931, underscoring this mitochondrial gene’s utility for resolving population structure in Southeast Asian mosquito vectors. Compared to COI (Hd=0.914, 35 haplotypes) and ITS2 (Hd=0.716, 5 haplotypes), COII shows intermediate resolution, reflecting concerted mtDNA evolution with elevated polymorphism suitable for phylogeographic inference [2]. Hap2 (n=6) predominates among Indonesian samples from East Nusa Tenggara, West Nusa Tenggara, and Central Sulawesi (positions 2, 4-5, 7, 12-13), indicating a core clade with regional connectivity. Hap9 (n=5) links Central Sulawesi, West Papua, West Java, and Lampung (positions 16-20), while Hap4 (n=2, North/Central Sulawesi at 6,10), and Hap5 (n=2, East Nusa Tenggara at 8-9) show localized clustering; singletons like Hap1 (West Sumatra), and Hap10 (Yogyakarta) highlight site-specific variants. Reference sequences from distinct groups: Hap11 (Australia), Hap12-17 (Thailand/China), suggesting geographic barriers or historical divergence from Indonesian populations [32]. Haplotype networks in this study would feature Hap2 as a central hub with short branches to Indonesian singletons (Hap 3,5-10), evidencing recent gene flow and star-like expansion across archipelago islands, while longer branches connect to Thailand/Chinese haplotypes (Hap12-17), implying isolation by distance or Pleistocene vicariance. This topology, with low reticulation, aligns with neutrality, indices signalling demographic growth, contrasting nuclear ITS2’s shallower structure, and supporting mtDNA’s maternal lineage tracking for vector dispersal models.

## 4. Discussion

### Genetic structuring and lineage differentiation in *Anopheles tessellatus* across Indonesia

This study provides a comprehensive molecular assessment of *An. tessellatus* populations across Indonesia using ITS2 nuclear and COI, COII mitochondrial markers, revealing clear evidence of genetic structuring and geographically associated lineages within what has traditionally been regarded as a single species. Although all analysed populations form a monophyletic group, phylogenetic reconstruction and genetic distance analyses consistently identify three principal lineages, broadly corresponding to Sumatra, Sulawesi, and Java–Nusa Tenggara. These findings support the hypothesis that *An. tessellatus* in Indonesia represents a genetically structured species complex undergoing intraspecific diversification, and possibly speciation due to geographic distances between these lineages. The concordant clustering patterns observed across nuclear and mitochondrial loci indicate that the detected structure reflects genuine evolutionary differentiation rather than stochastic variation or marker-specific bias. Similar multilocus congruence has been widely interpreted as evidence of lineage divergence in other *Anopheles* complexes, including *An. dirus*, *An. sundaicus*, and *An. maculipennis* [10, 30, 33].

### Comparative performance of ITS2, COI, and COII markers

The three genetic markers employed in this study exhibited complementary resolution. The nuclear ITS2 region showed moderate haplotype diversity and relatively low nucleotide divergence, consistent with the effects of concerted evolution acting on ribosomal DNA arrays [16]. Despite this evolutionary constraint, ITS2 successfully discriminated multiple population clusters, demonstrating its utility for identifying higher-level genetic structure within *An. tessellatus*.

In contrast, mitochondrial COI and COII genes revealed substantially higher haplotype diversity and finer-scale population differentiation. COI, in particular, displayed exceptional haplotype richness (Hd > 0.9), consistent with its established sensitivity to recent demographic processes and maternal lineage tracking in mosquito vectors [2, 34]. COII showed intermediate resolution, reinforcing its value as a complementary mitochondrial marker, as previously demonstrated in studies of *Anopheles superpictus* and *An. culicifacies* complexes [16, 17, 35]. The combined use of nuclear and mitochondrial markers, therefore, provides a robust framework for resolving population structure and detecting cryptic genetic diversity in *An. tessellatus*.

### Demographic history and haplotype structure

Across mitochondrial loci, Indonesian populations of *An. tessellatus* exhibits high haplotype diversity coupled with low nucleotide diversity, a genetic signature commonly associated with recent demographic expansion following historical bottlenecks [36]. Star-like haplotype networks observed in COI and COII analyses further support this interpretation, suggesting population growth from ancestral haplotypes with subsequent accumulation of localized mutations. Such demographic patterns have been reported in multiple mosquito species and are often linked to Pleistocene climatic oscillations, habitat expansion, or increased dispersal opportunities facilitated by anthropogenic environmental change [13, 37]. In Indonesia, repeated cycles of sea-level fluctuation during the Pleistocene likely created alternating phases of population isolation and connectivity, shaping the observed mitochondrial diversity.

### Biogeographic influences on lineage divergence

The geographic distribution of genetic lineages strongly reflects the complex biogeographic history of the Indonesian archipelago. The distinct clustering of Sulawesi populations across all markers is particularly notable and consistent with Sulawesi’s long-term tectonic isolation and composite geological origin [31, 38]. Wallacean barriers have been shown to restrict gene flow in numerous taxa, and similar patterns of lineage divergence have been documented in insects, molluscs, and vertebrates [39, 40]. Lineages associated with Sumatra and Java–Nusa Tenggara likely reflect historical fragmentation within Sundaland, combined with ecological heterogeneity and limited dispersal across marine barriers. Despite these isolating forces, genetic distances among Indonesian lineages remain below thresholds typically associated with fully reproductively isolated species, suggesting early-stage divergence rather than complete speciation.

### Implications for malaria and lymphatic filariasis control in Indonesia

From a public health perspective, the finding that *An. tessellatus* across Indonesia forms a genetically connected but regionally structured population has several implications. First, the presence of multiple nuclear and mitochondrial lineages suggests that local populations may differ in ecological preferences (e.g., larval habitats, attitude, microclimate), host choice, and behavior (indoor vs. outdoor biting, peak biting times), which can influence their capacity to sustain residual malaria and lymphatic filariasis transmission under current control measures [2, 41]. Second, high haplotype diversity raises the possibility that traits such as insecticide resistance, if they emerge, could spread regionally through movement of adults or passive transport, as observed in other *Anopheles* vectors [42].

Indonesia has committed to malaria elimination by 2030, and understanding the full spectrum of vector species and lineages, including secondary and potential vectors like *An. tessellatus,* is critical to achieving this goal [43, 44]. In several parts of South and Southeast Asia, *An. tessellatus* has been documented as a principal or secondary vector of malaria and lymphatic filariasis, suggesting that misidentification or underestimation of its role could compromise surveillance and intervention strategies [2, 41]. The genetic structure documented here provides a framework for prioritizing further entomological investigations, including blood-meal analysis, sporozoite detection, and insecticide susceptibility testing in key lineages (e.g., Sumatra and Sulawesi clusters) that exhibit elevated divergence and may represent functionally distinct vector populations.

### Limitations and future directions

Several limitations should be acknowledged. The sampling, while geographically extensive, remains uneven, with some islands and provinces represented by relatively few specimens, potentially underestimating local diversity and missing intermediate haplotypes. The study relies on three loci, one nuclear and two mitochondrial, which, although widely used and informative for species delimitation and population genetics in *Anopheles,* provide limited resolution compared to genome-wide single-nucleotide polymorphism or whole-genome approaches [19, 21, 34]. Furthermore, no ecological, behavioral, or vector competence data were integrated here, limiting the ability to link genetic lineages to epidemiological roles directly. Future research should therefore combine high-throughput genomic sequencing with targeted ecological and entomological investigations to test whether the genetic clusters identified in this study correspond to distinct eco-ethological units. Including additional nuclear markers or reduced-representation genomic data, such as RADseq or UCEs, would improve power to detect fine-scale structure and gene flow patterns, as demonstrated in other Southeast Asian taxa, such as squirrels and ricefish [40, 45]. Longitudinal sampling and integration of landscape and seascape features would further clarify how historical and contemporary barriers, such as deep ocean trenches and mountain ranges, shape the genetic architecture of *An. tessellatus* across Indonesia [46, 47].

## 5. Conclusions

This study demonstrates that *Anopheles tessellatus* populations in Indonesia are genetically structured into multiple geographically associated lineages, despite forming a single monophyletic group. Analyses of nuclear ITS2 and mitochondrial COI and COII markers consistently reveal high haplotype diversity, low nucleotide divergence, and limited gene flow among major island groups due to isolation by distance. These genetic patterns reflect the complex biogeographic history of the Indonesian archipelago and suggest ongoing intraspecific diversification within *An. tessellatus*. These findings highlight the presence of cryptic genetic structure in a medically important mosquito species and underscore the value of multilocus molecular approaches for elucidating evolutionary processes in vector populations.

Populations from West Sumatra, Central, and North Sulawesi, and Central Kalimantan exhibited genetic distances and unique haplotypes that may represent incipiently divergent lineages within the *An. tessellatus* complex and warrant an integrative taxonomic and ecological investigation. These aspects are particularly important to study considering the recognized and potential role of *An. tessellatus* as a malaria and lymphatic filariasis vector in Indonesia and neighbouring countries. These findings reinforce the need to incorporate this species complex into national vector surveillance programs, insecticide-resistance monitoring, and elimination planning efforts.

## Acknowledgments

We would like to express our gratitude to Lembaga Pengelola Dana Pendidikan (LPDP) or the Endowment Fund for Education Agency for financial support. The researcher extends gratitude to several teams and individuals who contributed to the study. These include the field team from Public Health Laboratory of Donggala (Risti, Ade Kurniawan, Arini Nur Syifa, Yulita Tribatha) for their assistance in mosquito collection in Central Sulawesi, and the Kulon Progo field team (Mujiyono, M. Choirul Hidajat, Muh. Fajri Rokhmad, Agung Puja Kesuma, Arief Mulyono, Mujiyanto, Rais Yunarko, Mara Ipa, Yusnita Mirna Anggraeny, Dian Eka Setyaningtyas, Tsabita Mutia, Rina Isnawati), Kulon Progo Health Office, Kali Bawang Community Health Center Team, team from Environmental Health Laboratory Center Salatiga (Aryo Ardanto, Valentinus Widi Ratno, Riyani Setiyaningsih) for their support. Additionally, acknowledgment is given to the laboratory and data analysis team, as well as all other parties who aided in conducting the research.

## Author contributions

Conceptualization: Anis Nurwidayati, Hari Purwanto, Raden Roro Upiek Ngesti Wibawaning Astuti, Budi Setiadi Daryono, Triwibowo Ambar Garjito, Sylvie Manguin.

Data curation: Anis Nurwidayati, Triwibowo Ambar Garjito, Lulus Susanti, Yuyun Srikandi.

Formal analysis: Anis Nurwidayati, Hari Purwanto, Raden Roro Upiek Ngesti Wibawaning Astuti,

Yudhi Ratna Nugraheni, Triwibowo Ambar Garjito, Sylvie Manguin.

Funding acquisition: Anis Nurwidayati, Sylvie Manguin.

Investigation: Anis Nurwidayati, Hari Purwanto, Raden Roro Upiek Ngesti Wibawaning Astuti, Lulus

Susanti, Yuyun Srikandi, Triwibowo Ambar Garjito.

Methodology: Anis Nurwidayati, Hari Purwanto, Raden Roro Upiek Ngesti Wibawaning Astuti, Budi

Setiadi Daryono, Triwibowo Ambar Garjito, Sylvie Manguin

Project administration: Anis Nurwidayati.

Resources: Anis Nurwidayati, Triwibowo Ambar Garjito, Lulus Susanti, Yuyun Srikandi, Sylvie Manguin.

Software: Anis Nurwidayati, Triwibowo Ambar Garjito, Sylvie Manguin.

Supervision: Hari Purwanto, Raden Roro Upiek Ngesti Wibawaning Astuti, Budi Setiadi Daryono,

Triwibowo Ambar Garjito, Sylvie Manguin.

Validation: Triwibowo Ambar Garjito, Sylvie Manguin.

Visualization: Anis Nurwidayati, Triwibowo Ambar Garjito, Sylvie Manguin.

Writing – original draft : Anis Nurwidayati

Writing – review & editing : Hari Purwanto, Raden Roro Upiek Ngesti Wibawaning Astuti,

Triwibowo Ambar Garjito, Sylvie Manguin.

